# Highly efficient *XIST* reactivation in female hPSC by transient dual inhibition of TP53 and DNA methylation during Cas9 mediated genome editing

**DOI:** 10.1101/2024.11.04.622001

**Authors:** Nami Motosugi, Keita Hasegawa, Natsumi Kurosaki, Erika Kawaguchi, Kenji Izumi, Yumi Iida, Misaki Higashiseto, Keiko Yokoyama, Ayumi Sasaki, Kazuhiko Nakabayashi, Atsushi Fukuda

## Abstract

Erosion of X-chromosome inactivation (XCI) complicates disease and developmental modeling in female human pluripotent stem cells (hPSCs). Previous studies demonstrated that Cas9-mediated editing of the *XIST* promoter *via* non-homologous end joining (NHEJ) or homology-directed repair (HDR) with a selection cassette upstream of *XIST* can trigger DNA demethylation and *XIST* reactivation, restoring XCI. Here, we show that NHEJ-mediated XCI reacquisition is more stable during differentiation than HDR. We further developed a novel, efficient *XIST* reactivation method by combining TP53 inhibition with DNA methylation maintenance suppression during Cas9-mediated NHEJ, offering a robust approach to achieving stable XCI in female hPSCs for diverse applications.

## Introduction

The irreversible silencing of the long non-coding RNA *XIST* in female human pluripotent stem cells (hPSCs) is largely unavoidable under conventional culture conditions (Anguera et al., 2012; Cloutier et al., 2022; Fukuda et al., 2021; Mekhoubad et al., 2012; Vallot et al., 2015). The loss of *XIST* expression leads to the erosion of X-chromosome inactivation (XCI), causing bi-allelic activation of X-linked genes and subsequent overexpression (Anguera et al., 2012; Mekhoubad et al., 2012; Motosugi et al., 2022). Given that the X-chromosome harbors over 500 genes (Sun et al., 2022), XCI erosion due to *XIST* silencing can significantly impact the outcomes of differentiation experiments in female hPSCs (Mekhoubad et al., 2012; Motosugi et al., 2022). In particular, disease modeling using female patient iPSCs, including those for X-linked gene disorders, has struggled to accurately recapitulate disease phenotypes *in vitro*. Thus, XCI erosion represents a major, often underappreciated, source of variability that compromises experimental reproducibility in studies utilizing female hPSCs.

Our previous work demonstrated that *XIST* repression in female hPSCs is caused by DNA methylation (DNAme) at *XIST* promoter regions, facilitated by *de novo* DNA methyltransferase activity (Fukuda et al., 2021). Recently, we showed that DNA hypermethylation could be reversed via Cas9-mediated non-homologous end joining (NHEJ), leading to *XIST* reactivation through endogenous transcriptional mechanisms (Motosugi et al., 2022). While DNA demethylation approach could reactivate *XIST* in female hPSCs, its efficiency was limited, with only ∼5% of transfected cells showing reactivation. To enhance this, we employed homology-directed repair (HDR) in Cas9-mediated gene editing, introducing a small targeting vector containing a 1.5 kb region of the *XIST* promoter along with an antibiotic selection marker. Following selection, *XIST* reactivation efficiency increased to ∼20% (Motosugi et al., 2022). However, the inclusion of exogenous sequences in up-stream of *XIST* gene during HDR raises concerns about potential unintended effects on *XIST* expression during disease modeling.

In this study, using iPSCs derived from a Rett syndrome (RTT) patient with an X-linked mental disorder (RTT-iPSCs) (Motosugi et al., 2022), we demonstrated that differentiating cells derived from RTT-hPSCs with NHEJ-mediated *XIST* reactivation could stably maintain *XIST* expression, highlighting that the approach can preserve integrity of dosage compensation. We further developed a highly efficient approach for *XIST* reactivation in female hPSCs by combining dual inhibition of TP53 and DNAme maintenance during Cas9-mediated NHEJ. This new highly efficient method for XCI de-erosion can improve the utility of various applications using female hPSCs.

### NHEJ mediated *XIST* reactivation maintains robust *XIST* expression in differentiating cell from female hPSCs

We previously established an iPSC line from a Rett syndrome (RTT) patient carrying the MECP2 mutation (R306H), in which *XIST* was reactivated using either NHEJ-mediated DNA demethylation or HDR-mediated renewal of the *XIST* promoter region with a selection cassette (RTT-iPSC^NHEJ^ and RTT-iPSC^HDR/CASS^, respectively) (Motosugi et al., 2022) (Fig. 1a). Using these RTT-iPSCs alongside healthy control lines (ADSC-iPSCs and SE1-5a^NHEJ^), all of which maintained *XIST* expression in approximately 80% cells (Motosugi et al., 2022), we generated cortical organoids to assess *XIST* and X-linked gene expression status during differentiation.

**Fig. 1.**
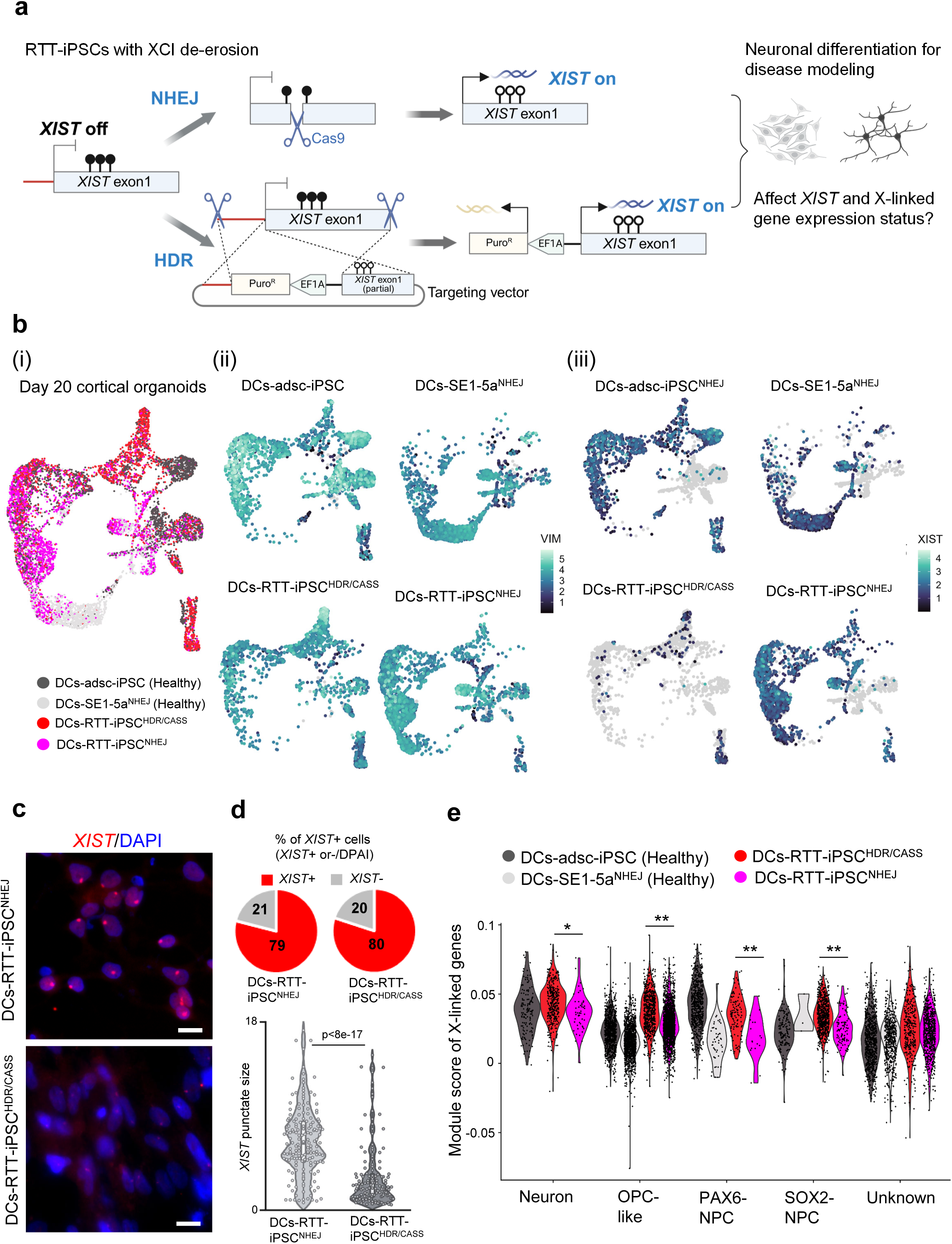
NHEJ mediated *XIST* reactivation maintains robust *XIST* expression in differentiating cell from female hPSCs. (**a**) Schematic overview of the experimental approach to assess *XIST* and X-linked gene expression in differentiating cells (DCs) derived from RTT-hPSCs. Two methods were used for *XIST* reactivation in RTT-iPSCs: (upper panel) DNA demethylation via Cas9-mediated non-homologous end joining (NHEJ), and (lower panel) homology-directed repair (HDR) using a gene targeting vector with an antibiotic selection cassette upstream of the *XIST* transcription start site (referred to as HDR/CASS). (**b**) scRNA-seq analysis of DCs. Cortical organoids at day 20 were subjected to scRNA-seq. (i-iii) Uniform manifold approximation and projection (UMAP) plots showing (i) cell line batch, (ii) *VIMENTIN* expression (an early neuronal marker), and (iii) *XIST* expression status (see also Fig. S1a). (**c**) Representative *XIST* RNA-FISH images in DCs derived from RTT-iPSCs at day 20 of cortical organoid differentiation. Scale bar: 20 μm. (**d**) Quantification of *XIST* RNA-FISH results. Upper panel: Percentage of *XIST*-positive cells; over 177 cells were analyzed in each group. Lower panel: *XIST* puncta size in DCs-RTT-iPSC^NHEJ^ and DCs-RTT-iPSC^HDR/CASS^, with each dot representing individual cell data. Statistical analysis was performed using t-tests. (**e**) X-linked gene module score analysis in DCs, compared across cell lines and cell types (see also Fig. S1b). Statistical analysis was performed using one-way ANOVA with post-hoc Tukey’s HSD test. *p < 0.01, **p < 0.001

On day 20 of differentiation, we performed single-cell RNA sequencing (scRNA-seq) to quantify *XIST* expression in differentiating cells (DCs). The scRNA-seq data showed that most cells were VIMENTIN-positive (+), indicating that the population was early neuronal differentiation phase (Eze et al., 2021) (Fig. 1b). Interestingly, over 50% of DCs derived from both healthy control and RTT-iPSC^NHEJ^ lines expressed *XIST*, while fewer than 10% of cells derived from the RTT-iPSC^HDR/CASS^ line were *XIST*+ (Fig. 1b and S1a). Notably, the small proportion of *XIST*+ cells in the RTT-iPSC^HDR/CASS^ line was enriched in neuronal clusters (Fig. S1b), consistent with previous findings showing a higher prevalence of *XIST*+ cells in neurons compared to other cell types within cortical organoids (Motosugi et al., 2022).

As the 10x Genomics 3’ expression system used in scRNA-seq captures only partial transcripts (Gao et al., 2020), we performed *XIST* RNA-FISH analysis on DCs-RTT-iPSC^NHEJ^ and DCs-RTT-iPSC^HDR/CASS^ lines using a probe targeting the entire *XIST* gene. FISH analysis at day 20 of differentiation confirmed that the fractions of *XIST*+ cells were similar between DCs-RTT-iPSC^NHEJ^ (79%) and DCs-RTT-iPSC^HDR/CASS^ (80%) (Fig. 1c and 1d). However, *XIST* foci in DCs-RTT-iPSC^HDR/CASS^ were significantly smaller than those in DCs-RTT-iPSC^NHEJ^ cells, consistent with the scRNA-seq findings (Fig. 1b-1d). These results indicated that *XIST* expression was reduced in DCs-RTT-iPSC^HDR/CASS^, suggesting that insertion of exogenous sequences like selection cassette might prevent stable *XIST* expression in DCs.

We next investigated whether *XIST* reduction in DCs-RTT-iPSC^HDR/CASS^ affected X-linked gene expression. Module score analysis of X-linked gene expression in scRNA-seq data revealed significantly lower scores for X-linked genes in DCs-RTT-iPSC^HDR/CASS^ compared to DCs-RTT-iPSC^NHEJ^, in both neuronal and non-neuronal clusters (Fig. 1e and S1b). Interestingly, RTT-derived DCs exhibited markedly higher scores in the OPC-like cluster compared to healthy control-derived DCs (Fig. 1e), suggesting that MECP2 mutations might severely affect XCI status in this cell cluster.

To further investigate molecular phenotypes in DCs derived from RTT-iPSCs, we conducted gene set enrichment analysis (GSEA) (Subramanian et al., 2005). Given the tendency of our differentiation protocol to favor OPC-like cells, we focused on both OPC-like clusters and neurons, a primary target cell type of RTT disease modeling. GSEA showed that certain pathways were differentially enriched in DCs-RTT-iPSC^NHEJ^ and DCs-RTT-iPSC^HDR/CASS^ (Supplementary Fig. S2). For example, epithelial-mesenchymal-transition (EMT) -related molecules were enriched in OPC-like clusters from DCs-RTT-iPSC^NHEJ^, though to a lesser extent in DCs-RTT-iPSC^HDR/CASS^ (Supplementary Fig. S2a). In neuron, IL6-Jak-Stat signaling was notably enriched in those derived from RTT-iPSC^NHEJ^ (Supplementary Fig. S2b). These findings suggested that *XIST* expression status influence molecular phenotypes in RTT disease modeling.

In conclusion, our data suggested that *XIST* expression induced *via* the NHEJ approach was stable after differentiation in both healthy and RTT-iPSC lines. While neuronal and glial differentiation was observed with both reactivation approaches, *XIST* repression and X-linked gene over-dosage in DCs-RTT-iPSC^HDR/CASS^ may lead to over-or under-estimation of disease phenotypes. These results highlighted the practical advantage of using NHEJ-mediated *XIST* reactivation to restore XCI in female hPSCs.

### Combining dual inhibition of TP53 and DNAme maintenance during Cas9-mediated NHEJ improves the efficiency on reactivating *XIST* in female hPSCs

We explored methods to enhance the efficiency of *XIST* reactivation through the NHEJ-mediated approach. Since transient inhibition of TP53 during Cas9-mediated gene editing—achieved by co-expressing a dominant-negative form of TP53—has been shown to significantly increase editing efficiency (Haapaniemi et al., 2018; Park et al., 2024), we investigated whether TP53 inhibition could similarly affect *XIST* reactivation. Additionally, as *XIST* reactivation via NHEJ was driven by DNA demethylation, we tested whether inhibiting DNMT1, essential for DNA methylation maintenance (Liao et al., 2015), could further improve reactivation efficiency.

Using ADSC-iPSCs lacking *XIST* expression, we first examined the effects of the DNMT1 inhibitor GSK3685032 on cell growth and *XIST* reactivation (Pappalardi et al., 2021). Continuous GSK3685032 treatment resulted in dose-dependent growth inhibition, with most cells dying after two weeks (Fig. S3a). These results were consistent with previous reports showing DNMT1’s role in hPSC survival (Liao et al., 2015). Despite poor growth, *XIST* was reactivated in a small subset of cells by GSK3685032 treatment (Fig. S3b), indicating that DNAme inhibition can promote *XIST* reactivation.

To further examine DNMT1 inhibition, we performed simple western analysis using 253G1-iPSCs (253G1), a well-characterized female hPSC line without *XIST* expression (Motosugi et al., 2021; Nakagawa et al., 2010). Treatment with 1 μM GSK3685032 significantly reduced DNMT1 protein levels within 24 hours (Fig. S3c), aligning with findings in mouse models (Chen et al., 2023).

Based on these findings, we evaluated the combined effects of TP53 and DNMT1 inhibition during Cas9-mediated NHEJ on *XIST* reactivation in 253G1 (Fig. 2a). For the DNAme inhibitor experiment, 253G1 were treated with 1 μM GSK3685032 for 24 hours before transfection and maintained in GSK3685032 for 48 hours post-transfection prior to EGFP sorting. As TP53 inhibition interferes with antibiotic selection such as puromycin (Jung et al., 2019), we used a Cas9-EGFP plasmid for transfection, followed by cell sorting based on EGFP expression (Fig. 2a). After sorting, cells were cultured, and XCI status was assessed by H3K27me3 immunofluorescence (IF), a hallmark of XCI (Fig. 2a). Inhibition of either DNAme or TP53 alone resulted in poor XCI reacquisition (<5% of cells), but dual inhibition significantly improved efficiency, increasing it to 28% (Fig. 2b).

**Fig. 2.**
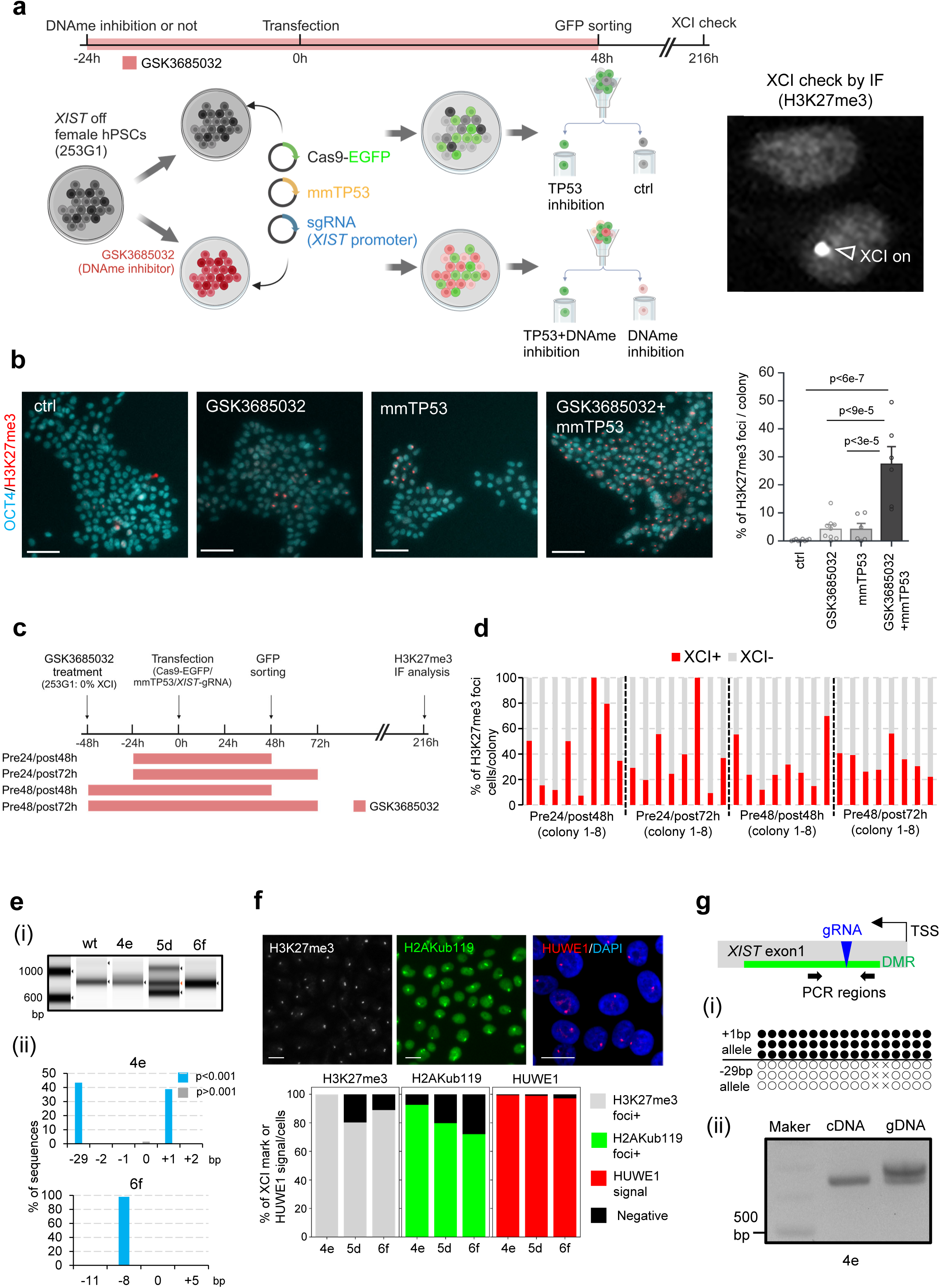
Combining dual inhibition of TP53 and DNAme maintenance during Cas9-mediated NHEJ improves the efficiency on reactivating *XIST* in female hPSCs. (**a**) Experimental scheme for assessing the effects of TP53 and DNAme inhibition on *XIST* reactivation in female hPSCs. For DNAme inhibition, the DNMT1-specific inhibitor GSK3685032 was applied. Cells were pre-treated with (for 24 hours) or without GSK3685032 before transfection, and those in the DNAme inhibition group were maintained in GSK3685032 for 48 hours post-transfection prior to GFP sorting. Sorted cells were then assessed for H3K27me3 foci (XCI hallmark) by immunofluorescence (IF). The representative image on the right shows H3K27me3 foci. In transfection, a vector expressing a dominant-negative form of TP53 (mmTP53) was co-transfected with Cas9-EGFP and gRNA vectors targeting the *XIST* promoter region. (**b**) Representative images of H3K27me3 and OCT4 staining, along with quantification results. Each dot represents the percentage of H3K27me3 foci cells with OCT4 expression within individual colonies, with at least 218 cells analyzed per colony. Statistical analysis was conducted using one-way ANOVA with post-hoc Tukey’s HSD test. Scale bar: 100 μm. (**c**) Experimental scheme for optimizing the timing of GSK3685032 treatment. (**d**) Quantitative results showing the percentage of H3K27me3 foci-positive cells within colonies. Only OCT4-positive cells were analyzed, with at least 383 cells counted per colony. (**e**) Genotyping analysis of sub-cloned *XIST*-reactivated 253G1-derived lines (4e, 5d, and 6f). (i) Automated electrophoresis image and (ii) TIDE analysis for genomic scar evaluation in lines 4e and 6f. (**f**) XCI status assessment. (Upper panel) Representative images of H3K27me3, H2AKub119 foci, and HUWE1 expression status. IF was used for H3K27me3 and H2AKub119 analysis, and RNA-FISH was used for HUWE1 expression. In FISH assay, DNA was stained with DAPI. Scale bar: 20 μm. (Lower panel) Quantification results based on analysis of over 316 cells in IF and 139 cells in FISH per line. (**g**) DNAme and allelic expression analysis at the *XIST* locus. Schematic of gRNA, PCR primer, and differentially methylated regions (DMRs) in *XIST* exon 1. (i) Bisulfite sequencing analysis and (ii) allelic expression assessment for the 4e line.

Next, we optimized GSK3685032 treatment to further enhance XCI reacquisition. Cells were treated with GSK3685032 for 24–48 hours before transfection, with an additional 24-hour treatment post-sorting (72h post-transfection) (Fig. 2c). After night days of culture, IF analysis for H3K27me3 and OCT4 showed that 32.1% to 43.7% of cells were XCI-positive with pluripotency (Fig. 2d). Notably, in the pre48/post72h condition, over 22% of cells across all colonies were H3K27me3-positive (Fig. 2c and 2d), demonstrating the efficacy of prolonged GSK3685032 treatment for XCI reacquisition.

We further isolated three subclones with >80% H3K27me3-positive cells (termed as 4e, 5d, and 6f lines) (Fig. 2e). Genotyping analysis revealed NHEJ-induced indels, consistent with previous findings (Fig.2e). To verify whether XCI reacquisition occurred on the previously eroded X chromosome, we performed RNA-FISH for HUWE1, a gene retaining its inactive status post-XCI erosion (Motosugi et al., 2022; Patel et al., 2017). HUWE1 was detected in 97% of cells, confirming XCI reacquisition on the eroded chromosome (Fig. 2f). Additional IF analysis for H2AKub119, another XCI marker, showed foci in >70% of OCT4-positive cells (Fig. 2f), further supporting successful XCI restoration.

Next, to examine the relationship between DNAme at the *XIST* promoter and its expression, we performed bisulfite sequencing and allele-specific expression analysis on the 4e line, which harbored a 29-bp deletion (Fig. 2e). Bisulfite sequencing showed hypomethylation of the *XIST* promoter on the deleted allele (Fig. 2g), and *XIST* expression was confirmed from this allele (Fig. 2g), consistent with previous reports (Motosugi et al., 2022).

In conclusion, our results demonstrated that dual inhibition of TP53 and DNMT1 significantly enhanced the efficiency of XCI reacquisition in female hPSCs.

### Scalable expansion and reproducibility of XCI-restored female hPSCs *via* dual inhibition

Our previous study demonstrated that *XIST* reactivation in female hPSCs is not maintained during long-term culture (Motosugi et al., 2022), although this process could be slowed by the continuous use of a Rho-kinase inhibitor (ROCKi) in the culture medium. To build on this, we investigated whether ROCKi could support the scalable expansion of XCI-restored lines (4e, 5d, and 6f) while maintaining XCI. We seeded three XCI-restored clones at a density of 2.5 x 10⁶ cells and expanded them to over 1 x 10⁷ cells (Fig. 3a). IF analysis for H3K27me3 and OCT4 confirmed that 69–95% of cells retained XCI along with pluripotency markers (Fig. 3b), demonstrating that dual inhibition-mediated XCI restoration enables scalable expansion.

**Fig. 3.**
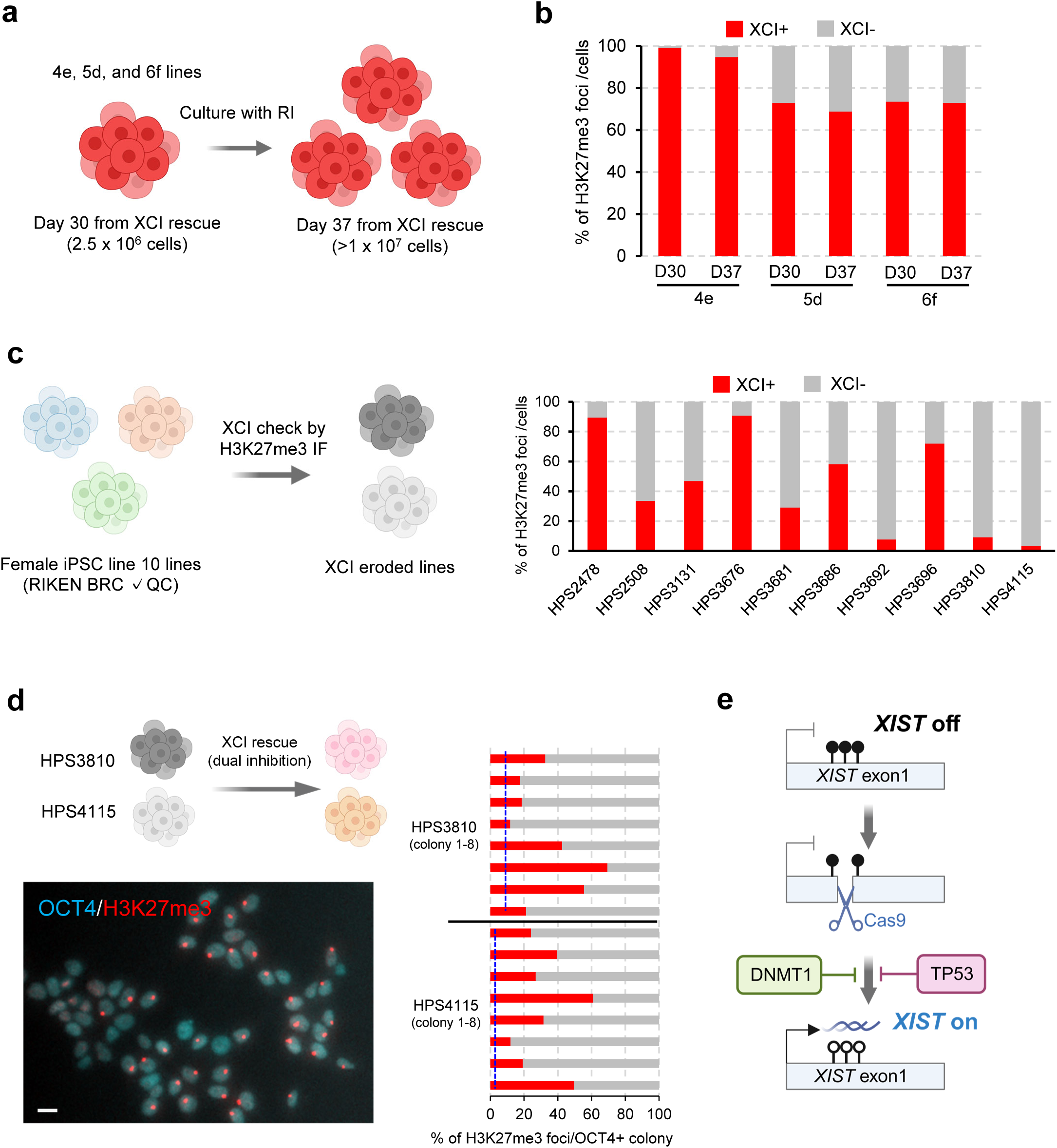
Scalable expansion and reproducibility of XCI-restored female hPSCs via dual inhibition approach. (**a**) Schematic of scalable expansion of XCI-rescued lines. Day 0 represents the day of transfection. XCI-rescued lines were cultured to reach over 10⁷ cells in the presence of RI and assessed for XCI status by IF of H3K27me3 and OCT4. (**b**) Quantification of XCI-positive cells. At least 584 cells were analyzed per line to confirm XCI restoration. (**c**) XCI status assessment in 10 RIKEN BRC quality-controlled (QC) female iPSC lines. The right panel shows quantification of XCI status by IF for H3K27me3, restricted to OCT4-positive cells. At least 382 cells were analyzed per line. (**d**) XCI rescue in HPS3810 and HPS4115 lines. The representative image of H3K27me3 foci with OCT4 staining (Left lower panel). The right panel shows the percentage of cells with H3K27me3 foci within the OCT4-positive population before sub-cloning. The blue dotted line represents the starting population shown in (c). Scale bar: 20 μm. (**e**) Model of dual inhibition approach for *XIST* reactivation. Transient inhibition of TP53 and DNMT1 during the NHEJ process boosts genome editing efficiency and prevents DNA methylation maintenance, thereby enhancing the efficiency of *XIST* reactivation.

Next, we assessed the reproducibility of this dual inhibition approach in other female hPSCs. We used healthy female hiPSC lines from the RIKEN BioResource Center (BRC), which were quality-controlled (QC) and available for global distribution. We examined the XCI status of 10 QC-verified female hiPSC lines *via* H3K27me3 staining. Of these, three lines retained XCI in over 70% of cells, while the others exhibited varying degrees of XCI erosion, ranging from 3% to 91% (Fig. 3d). We selected the 4115 and 3810-iPSC lines, which exhibited largely eroded XCI status (H3K27me3 foci present in fewer than 10% of cells), to test the reproducibility of dual inhibition-mediated XCI reacquisition. These iPSC lines were treated with GSK3685032 under the pre48/post72h condition, with TP53 inhibition during transfection, as shown in Fig. 2c. Following EGFP sorting, the cells were cultured and subjected to IF analysis for H3K27me3 and OCT4. Approximately 33% of cells in the colonies before sub-cloning displayed XCI reacquisition with OCT4 positivity in both lines (Fig. 3e), demonstrating that the dual inhibition approach was effective in restoring XCI in additional female iPSC lines (Fig. 3f).

## Discussion

Since GSK3685032 has been identified as a specific inhibitor of DNMT1 (Chen et al., 2023; Pappalardi et al., 2021), our findings indicated that DNA methylation (DNAme) following Cas9-induced double-strand breaks (DSBs) was maintained by DNMT1, rather than de novo DNA methyltransferases (Fukuda et al., 2021; Motosugi et al., 2022). Once established, DNAme is stably preserved by DNMT1 in various genomic regions, including imprinted genes. Previous studies have shown that NHEJ-mediated gene reactivation via DNA demethylation was successfully applied to the imprinted gene GTL2 (Motosugi et al., 2022). Therefore, the dual inhibition approach would induce the efficient reactivation of genes regulated by DNAme. However, considering that NHEJ often introduces genomic indels, potentially leading to aberrant protein expression in coding genes, this approach may be more appropriate for non-coding RNAs.

From a biological perspective, we observed that *XIST* expression via the HDR approach was significantly reduced in DCs derived from RTT-iPSCs (Fig. 1b-1e). Combining *XIST* reduction with small molecules has been proposed as a strategy to reactivate the wild-type MECP2 allele for RTT treatment (Grimm and Lee, 2022). However, GSEA in DCs-RTT-iPSCs indicated that *XIST* reduction might also help mitigate aberrant activation of pathways like TNF-α signaling in neurons (Supplementary Fig. S2b). Nevertheless, given the opposing effects observed in pathways such as hedgehog signaling, careful consideration is needed when applying *XIST* reduction strategies for therapeutic purposes.

In both disease and developmental modeling, maintaining *XIST* expression in female cells is crucial for accurately interpreting cellular phenotypes. Many studies utilizing female hPSCs have relied on stable lines, such as 253G1 used in this study, which often exhibit XCI erosion. The erosion is particularly problematic when patient-specific iPSCs are employed for genotype-phenotype studies. However, the dual inhibition approach can efficiently reactivate *XIST* expression in female hPSCs, offering a valuable tool for enhancing the accuracy and utility of patient-specific iPSC models.

## Methods

All human ES cell lines in this study were used under the instruction on the use of human embryonic stem cell research guidelines of Tokai University, the National Center for Child Health and Development, and the Ministry of Education in Japan. The experiments on X-linked disease iPSCs were approved by the Institutional Review Board for Clinical Research, Tokai University.

### hPSC culture

Human pluripotent stem cells (hPSCs) were maintained in StemFlex medium (Thermo Fisher Scientific, Waltham, MA, USA) on plates coated with hES cell-grade Matrigel (Corning Inc., Corning, NY, USA). For the passaging of hPSCs, the culture medium was supplemented with 1 mM EDTA and 10 μM Y-27632 (StemCell Technologies, Vancouver, Canada), and cells were allowed to incubate for 24 hours, unless indicated otherwise. All hPSCs were cultured as described above and at 37°C in the presence of 5% CO2. XIST reactivated RTT-iPSCs (HPS3049), ADSC-iPSC, SE1-5a, and 253G1 lines were previously reported (REF). The ten quality-controlled (QC) female hiPSC lines (HPS2478, HPS2508, HPS3131, HPS3676, HPS3681, HPS3686, HPS3692, HPS3696, HPS3810, HPS4115) were obtained from the RIKEN BioResource Center (https://cell.brc.riken.jp/ja/).

### *XIST* reactivation via dual inhibition approach

hPSCs were treated with GSK3685032 (#HY-139664; MedChemExpress, NJ, USA) for indicated periods and concentrations in each figure. These hPSCs were dissociated into single cell using TrypLE Express (#12605010; Thermo Fisher Scientific) and subjected to transfection (Neon Transfection System, Thermo Fisher Scientific) of Cas9-T2A-EGFP-ires-puro (a gift from Timo Otonkoski, #78311; Addgene), pCE-mp53DD (a gift from Shinya Yamanaka Plasmid #41856), pSPgRNA (a gift from Charles Gersbach, #47108; Addgene), targeting *XIST* promoter (REF). after transfection, hPSCs were cultured in the presence of GSK3685032 or not for two days followed by FACS sorting (BD, Franklin Lakes, NJ, USA). EGFP+ or – cells were sorted and cultured until further experiments. For sub-cloning of XIST reactivated cells, each colony was picked up and plated into 96 well plate. After 70-80% of confluent, the cells were splitted into 2 replicates: one was used for H3K27me3 IF stating for screening of the cells with XCI, the other was used for further experiments.

### Generation of cortical organoids

Differentiation into cortical organoids was conducted as previously reported. In brief, cortical organoids were generated from hPSCs using the STEMdiff Cerebral Organoid Kit (#08570; StemCell Technologies) following the manufacturer’s protocol. hPSCs were dissociated into single cells with TrypLE Express (Thermo Fisher Scientific), and 9,000 cells were seeded into a 96-well ultra-low attachment plate (#7007; Corning Inc.) to form embryoid bodies (EBs) over 5 days. EBs were then transferred to a 24-well ultra-low attachment plate (#3473; Corning Inc.) in neural induction medium for 48 hours before being embedded in Matrigel. After polymerization at 37°C, expansion medium was added. On day 10, maturation medium was introduced, and the cultures were placed on an orbital shaker with medium changes every 3–4 days until further analysis.

### Immunofluorescence analysis

Immunofluorescence (IF) analysis was performed as previously described (Fukuda et al., 2021; Motosugi et al., 2021). Cells were fixed with 4% paraformaldehyde (PFA) in PBS for 20 minutes, permeabilized with 0.1% Triton-X for 20 minutes, and blocked with 1.5% BSA in PBS for 1 hour. Primary and secondary antibody reactions were performed sequentially, followed by imaging using an Axio Imager 2 microscope (Carl Zeiss) or Keyence BZ-9000 system (Keyence). Antibody details are provided in Table S1.

### RNA-FISH

RNA-FISH was performed following previously established protocols (Fukuda et al., 2021; Motosugi et al., 2021). Cells were fixed with 4% PFA in PBS for 20 minutes, permeabilized with 0.1% Triton-X for 20 minutes, and subjected to RNA-FISH. Probes for TRE-hXIST (REF) and HUWE1 (BAC clones CTD 3063K22, PR11-975N19) were prepared using a nick translation kit (Abbott Laboratories). VECTASHIELD with DAPI was used for nuclear staining. Images were captured using an Axio Imager 2 microscope (Carl Zeiss). For organoid analysis, organoids derived from RTT-iPSCs were dissociated with the Neural Tissue Dissociation Kit (P) (Miltenyi Biotec), plated on poly-d-lysine-coated coverslips, and processed for RNA-FISH as described.

### Genotyping and PCR

*XIST* reactivated hPSCs were subjected to genotyping PCR using primer spanning gRNA targeting regions and checked by TapeStation (Agilent, Santa Clara, CA, USA) or electrophoresis. To check for scars in each gene editing, TIDE (http://shinyapps.datacurators.nl/tide/) analysis was performed around the gRNA targeting sequences. All primer sequences are listed in Table S1.

### Simple western analysis

Capillary western analyses were performed using the ProteinSimple Wes System according to manufacture’s instruction. Briefly, proteins from hPSCs were extracted using RIPA lysis and extraction buffer (Thermo Fisher Scientific). Protein concentrations were quantified and analyzed for a total of 2.4 ug of each sample. Anti-DNMT1 (ab188453, Abcam) was used for the assay. The data was analyzed using Compass software (ProteinSimple).

### Bisulfite sequencing

Bisulfite sequencing was performed as previously described (Fukuda et al., 2021). Genomic DNA was extracted using the EZ DNA Methylation Kit (Zymo Research). Amplified DNA was cloned into the pGEM-T Easy Vector (Promega) and sequenced. Data were analyzed using QUMA (http://quma.cdb.riken.jp/). Primer sequences are provided in Table SX.

### Single-cell RNA-sequencing analysis

Single-cell RNA sequencing was performed using the 10× Chromium platform (10× Genomics, Chromium Single Cell 3’ Reagent Kits) at Genewiz (South Plainfield, NJ, USA) following the manufacturer’s protocol. Data alignment to the GRCh38 human genome was conducted with Cell Ranger (v3.1.0, 10× Genomics), and filtered gene expression matrices were generated using default settings. Data were analyzed in R using the Seurat package (v5) (Hao et al., 2024). Cells with more than 200 unique features and less than 25% mitochondrial content were retained for analysis. UMAP dimensionality reduction was applied using the “integrated.mnn” method, and cell type annotation was conducted using the FindMarkers function. Module score analysis was performed on X-linked genes, excluding *XIST*, using the AddModuleScore function. The raw and processed data are deposited in GEO (GSE280890 * will not be released until accepted).

### Quantification and statistical analysis

Statistical analyses for the scRNA-seq data were conducted using the Seurat package. For multiple comparisons among different samples were evaluated using ANOVA followed by post-hoc tests, implemented in R. For other assays, including XCI+ percentage, paired two-tailed t-tests or ANOVA followed by post-hoc tests were performed in R, with p-values < 0.05 considered statistically significant. The number of cells analyzed are provided in the figure legends.

## Acknowledgements

We would like to thank the members of the Support Center for Medical Research and Education, Tokai University School of Medicine, for helping with the various experiments. The study was supported by AMED under Grant Number 22bm0804030h0002 to A.F.

## Author contributions

A.F. conceived the project and designed the experiments. N.M. K.H., N.K., E.K., K.I., Y.I., M.H., K.Y., and A.S. conducted the experiments. N.M., K.H., N.K., E.K., K.I., K.N., and A.F. analyzed the data. All authors interpreted the data and A.F. supervised the project. A.F. wrote the manuscript with input from all authors.

## Competing interests

A.F. is applying a patent for the method to generate *XIST* re-expressing female hPSCs by gRNA-mediated *XIST* demethylation (2022-062974).

**Supplementary Fig. S1.** scRNA-seq analysis of DCs derived from RTT-iPSCs with *XIST* reactivation. (a) *XIST* expression levels across all analyzed cells. (b) UMAP plots showing (i) cell type annotations and (ii–v) expression levels of key cell-type markers: (ii) SOX2 and (iii) PAX6 for neuronal progenitors, (iv) PDGFRA for OPCs, and (v) TUBB3 for neurons.

**Supplementary Fig. S2.** GSEA analysis in DC clusters. (a, b) GSEA of the OPC-like cluster (a) and neuronal cluster (b). Notably, no neuronal cluster was detected in the DCs-SE1-5a^NHEJ^ line (see also Supplementary Fig. S1b v).

**Supplementary Fig. S3.** Effects of GSK3685032 on female hPSC growth and DNMT1 expression. (a) Representative bright-field image of GSK3685032-treated 253G1 cells; scale bar: 500 μm. (b) *XIST* RNA-FISH analysis. The representative images of GSK3685032-treated 253G1 cells with DNA stained by DAPI; scale bar: 20 μm. (c) Wes analysis of DNMT1 protein levels. Left: chromatographic profiles for each treatment group. Right: quantification of DNMT1 expression. Statistical analysis performed using t-tests; *P < 0.0001.

## References

1. Anguera, M.C., Sadreyev, R., Zhang, Z., Szanto, A., Payer, B., Sheridan, S.D., Kwok, S., Haggarty, S.J., Sur, M., Alvarez, J., et al. (2012). Molecular signatures of human induced pluripotent stem cells highlight sex differences and cancer genes. Cell Stem Cell 11, 75–90.

2. Chen, Q., Liu, B., Zeng, Y., Hwang, J.W., Dai, N., Correa, I.R., Jr., Estecio, M.R., Zhang, X., Santos, M.A., Chen, T., et al. (2023). GSK-3484862 targets DNMT1 for degradation in cells. NAR Cancer 5, zcad022.

3. Cloutier, M., Kumar, S., Buttigieg, E., Keller, L., Lee, B., Williams, A., Mojica-Perez, S., Erliandri, I., Rocha, A.M.D., Cadigan, K., et al. (2022). Preventing erosion of X-chromosome inactivation in human embryonic stem cells. Nat Commun 13, 2516.

4. Eze, U.C., Bhaduri, A., Haeussler, M., Nowakowski, T.J., and Kriegstein, A.R. (2021). Single-cell atlas of early human brain development highlights heterogeneity of human neuroepithelial cells and early radial glia. Nat Neurosci 24, 584–594.

5. Fukuda, A., Hazelbaker, D.Z., Motosugi, N., Hao, J., Limone, F., Beccard, A., Mazzucato, P., Messana, A., Okada, C., San Juan, I.G., et al. (2021). De novo DNA methyltransferases DNMT3A and DNMT3B are essential for XIST silencing for erosion of dosage compensation in pluripotent stem cells. Stem Cell Reports 16, 2138–2148.

6. Gao, C., Zhang, M., and Chen, L. (2020). The Comparison of Two Single-cell Sequencing Platforms: BD Rhapsody and 10x Genomics Chromium. Curr Genomics 21, 602–609.

7. Grimm, N.B., and Lee, J.T. (2022). Selective Xi reactivation and alternative methods to restore MECP2 function in Rett syndrome. Trends Genet 38, 920–943.

8. Haapaniemi, E., Botla, S., Persson, J., Schmierer, B., and Taipale, J. (2018). CRISPR-Cas9 genome editing induces a p53-mediated DNA damage response. Nat Med 24, 927–930.

9. Hao, Y., Stuart, T., Kowalski, M.H., Choudhary, S., Hoffman, P., Hartman, A., Srivastava, A., Molla, G., Madad, S., Fernandez-Granda, C., et al. (2024). Dictionary learning for integrative, multimodal and scalable single-cell analysis. Nat Biotechnol 42, 293–304.

10. Jung, J.H., Lee, H., Kim, J.H., Sim, D.Y., Ahn, H., Kim, B., Chang, S., and Kim, S.H. (2019). p53-Dependent Apoptotic Effect of Puromycin via Binding of Ribosomal Protein L5 and L11 to MDM2 and its Combination Effect with RITA or Doxorubicin. Cancers (Basel) 11.

11. Liao, J., Karnik, R., Gu, H., Ziller, M.J., Clement, K., Tsankov, A.M., Akopian, V., Gifford, C.A., Donaghey, J., Galonska, C., et al. (2015). Targeted disruption of DNMT1, DNMT3A and DNMT3B in human embryonic stem cells. Nat Genet 47, 469–478.

12. Mekhoubad, S., Bock, C., de Boer, A.S., Kiskinis, E., Meissner, A., and Eggan, K. (2012). Erosion of dosage compensation impacts human iPSC disease modeling. Cell Stem Cell 10, 595–609.

13. Motosugi, N., Okada, C., Sugiyama, A., Kawasaki, T., Kimura, M., Shiina, T., Umezawa, A., Akutsu, H., and Fukuda, A. (2021). Deletion of lncRNA XACT does not change expression dosage of X-linked genes, but affects differentiation potential in hPSCs. Cell Rep 35, 109222.

14. Motosugi, N., Sugiyama, A., Okada, C., Otomo, A., Umezawa, A., Akutsu, H., Hadano, S., and Fukuda, A. (2022). De-erosion of X chromosome dosage compensation by the editing of XIST regulatory regions restores the differentiation potential in hPSCs. Cell Rep Methods 2, 100352.

15. Nakagawa, M., Takizawa, N., Narita, M., Ichisaka, T., and Yamanaka, S. (2010). Promotion of direct reprogramming by transformation-deficient Myc. Proc Natl Acad Sci U S A 107, 14152–14157.

16. Pappalardi, M.B., Keenan, K., Cockerill, M., Kellner, W.A., Stowell, A., Sherk, C., Wong, K., Pathuri, S., Briand, J., Steidel, M., et al. (2021). Discovery of a first-in-class reversible DNMT1-selective inhibitor with improved tolerability and efficacy in acute myeloid leukemia. Nat Cancer 2, 1002–1017.

17. Park, J.C., Kim, Y.J., Hwang, G.H., Kang, C.Y., Bae, S., and Cha, H.J. (2024). Enhancing genome editing in hPSCs through dual inhibition of DNA damage response and repair pathways. Nat Commun 15, 4002.

18. Patel, S., Bonora, G., Sahakyan, A., Kim, R., Chronis, C., Langerman, J., Fitz-Gibbon, S., Rubbi, L., Skelton, R.J.P., Ardehali, R., et al. (2017). Human Embryonic Stem Cells Do Not Change Their X Inactivation Status during Differentiation. Cell Rep 18, 54–67.

19. Subramanian, A., Tamayo, P., Mootha, V.K., Mukherjee, S., Ebert, B.L., Gillette, M.A., Paulovich, A., Pomeroy, S.L., Golub, T.R., Lander, E.S., et al. (2005). Gene set enrichment analysis: a knowledge-based approach for interpreting genome-wide expression profiles. Proc Natl Acad Sci U S A 102, 15545–15550.

20. Sun, Z., Fan, J., and Wang, Y. (2022). X-Chromosome Inactivation and Related Diseases. Genet Res (Camb) 2022, 1391807.

21. Vallot, C., Ouimette, J.F., Makhlouf, M., Feraud, O., Pontis, J., Come, J., Martinat, C., Bennaceur-Griscelli, A., Lalande, M., and Rougeulle, C. (2015). Erosion of X Chromosome Inactivation in Human Pluripotent Cells Initiates with XACT Coating and Depends on a Specific Heterochromatin Landscape. Cell Stem Cell 16, 533–546.

