## Supplemental Figure 1, 2, 3, and Supplemental Table 1 for "Highly efficient *XIST* reactivation in female hPSC by transient dual inhibition of TP53 and DNA methylation during Cas9 mediated genome editing"

### *Fig. S1. scRNA-seq analysis in DCs derived from RTT-iPSCs with XIST reactivation*

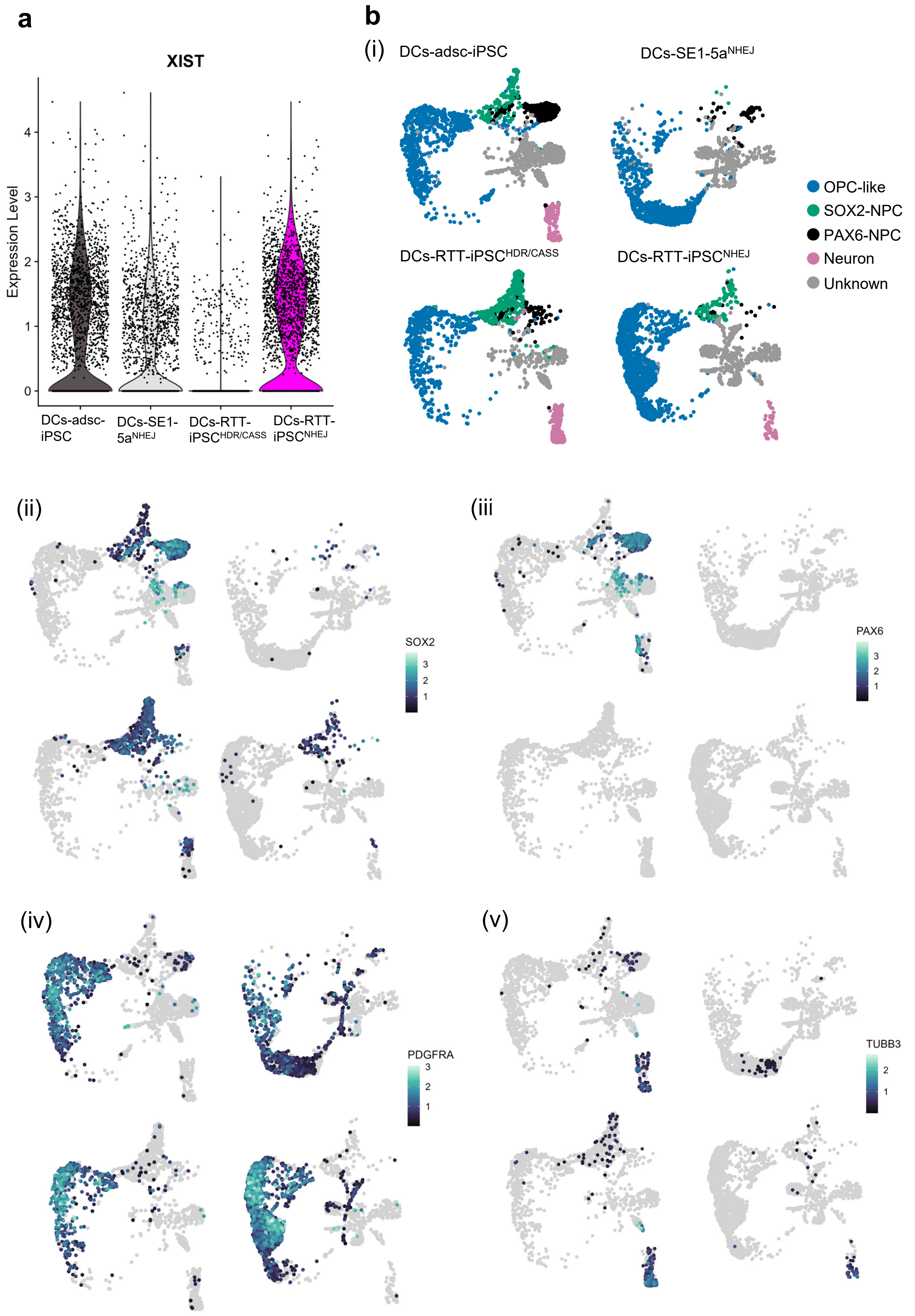

### *Fig. S2. GSEA analysis in DCs*

**a**

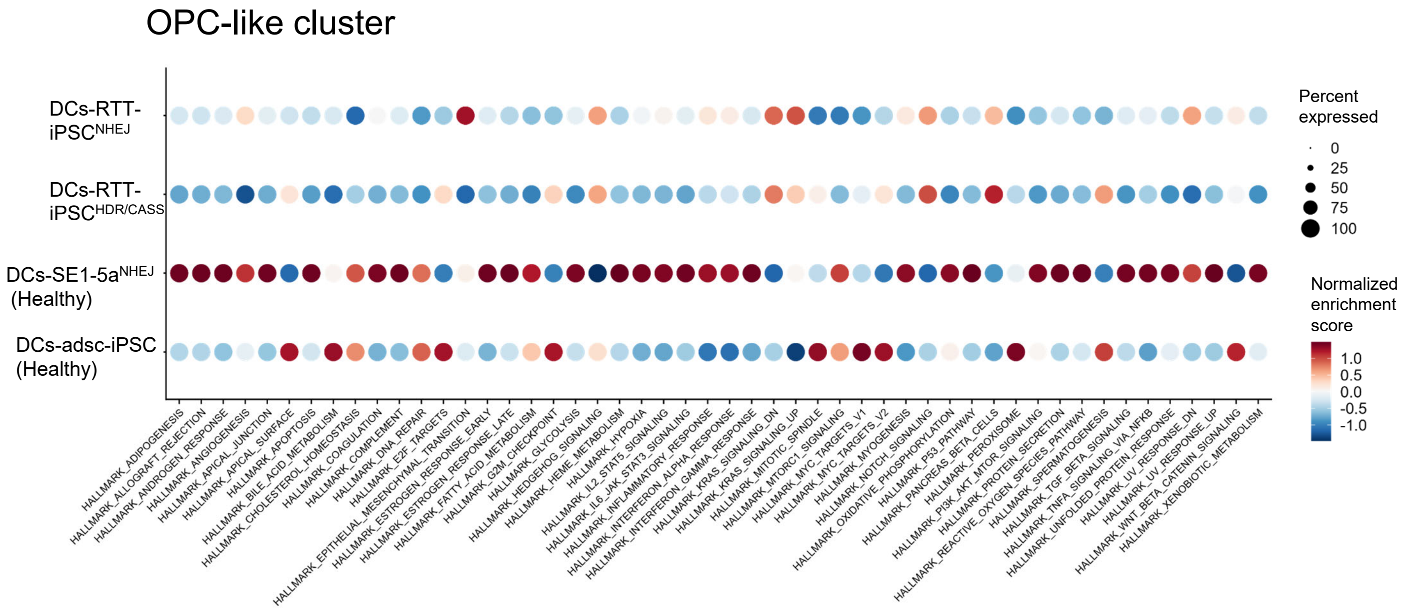


**b**

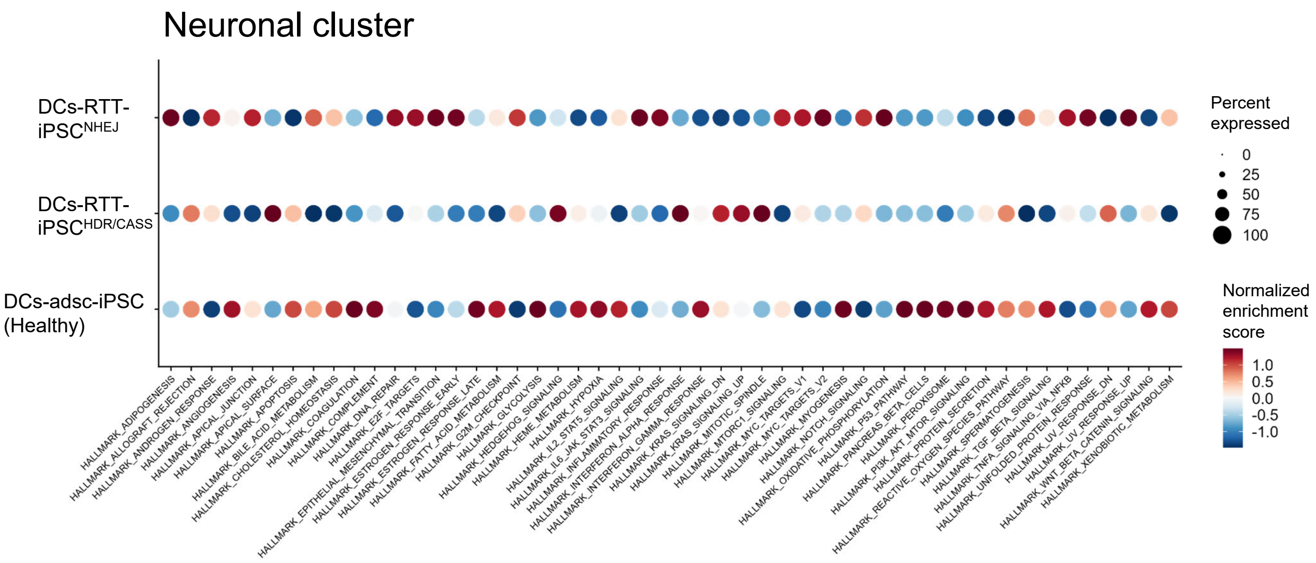

***Fig. S3. Effect of GSK3685032 on female hPSC growth and DNMT1 expression status***

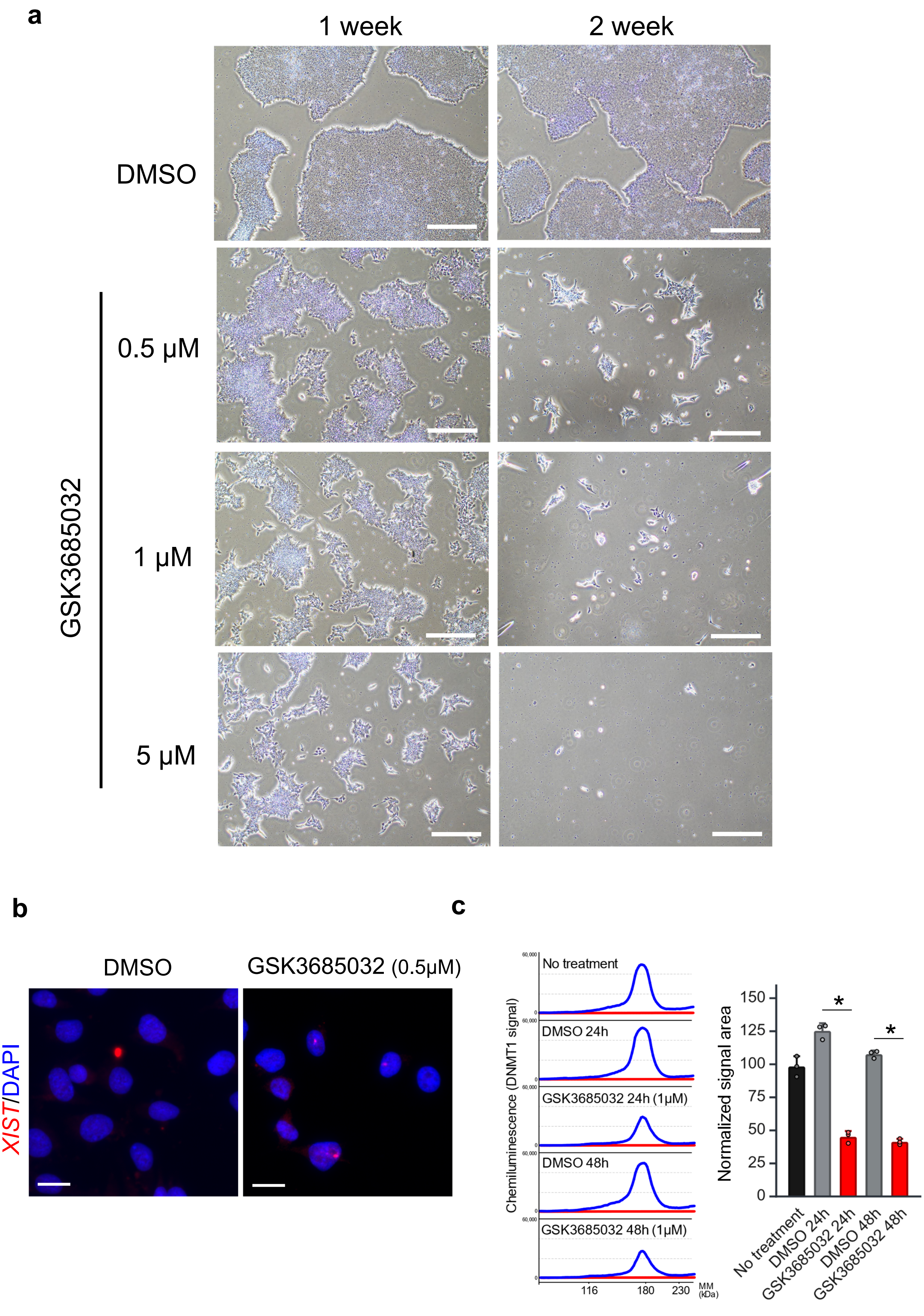

**Table S1. Primer and antibody information**

| Primer | Sequences | assay |
|---|---|---|
| 5_region_F2 | TTCTCTGCCAAAGCGGTAGGTACACT | TIDE |
| 5_region_R1 | CTCCTAGTGTCTTCTTGACACGTCC | |
| BS_test_6-F | GTGTAGATATTTTAGAGAGTGTAATAATTT | BS |
| BS_test_6-R | AAATACCTACCTTTTAATTCTTTTTTATTC | |

| Antibody name | Manufacture | Catalog No. | Dilution ratio | assay |
|---|---|---|---|---|
| H3K27me3 Rabbit monoclonal | Cell Signaling | 9733 | 1:500 | IF |
| OCT4 Mouse monoclonal | Santa Cruz | sc-5279 | 1:500 | |
| H2AKub119 Rabbit monoclonal | Cell Signaling | 8240 | 1:200 | |
| DNMT1 | Abcam | ab188453 | 1:250 | WES |
